# FMSClusterFinder: A new tool for detection and identification of clusters of sequential motifs with varying characteristics inside genomic sequences

**DOI:** 10.1101/2022.01.23.474238

**Authors:** Mohammad Mahdi Hejazi, Faegheh Golabi, Mohsen Bahrami, Houman Kahroba, Mohammad Saeid Hejazi

## Abstract

This paper describes FMSClusterFinder, a new tool and algorithm for identification and detection of clusters of sequential blocks inside the DNA and RNA subject sequences. Gene expression and genomic groups’ performance is under the control of functional elements cooperating with each other as clusters. The functional motifs or blocks are often comparably short, degenerate and are located within varying distances from each other. Since functional motifs mostly act in relation to each other as clusters, finding such clusters of blocks is an effective approach to identify functional groups and their function and structure, which represents the need for development of new corresponding tools.

The presented web application finds clusters of sequential blocks, with even altering sequences and located in varying distances from each other inside the subject sequences, simultaneously. Additionally, the blocks could be searched with user defined constant or varying characteristics such as: a) different levels of similarity, b) varying minimum number of blocks required to build up the query cluster, c) different types of sequence (degenerate or standard) and d) one or multiple alternative sequences for each block. FMSClusterFinder is available freely at http://fmsclusterfinder.fmsbiog.com.

## Introduction

The functional cis-acting elements build up the genome performance basis, which is raised from interaction of cis and trans-acting groups. Accordingly, predicting functional features of the genome beyond protein coding regions has received increasing attention as sequencing of genomes becomes widespread (Ian Dunham et al. 2012; Yue et al. 2014). The discovery of binding patterns in DNA, RNA and protein has led to determination of many functional sites such as open reading frames, promoter elements of genes, enhancers, silencers, intron/exon splicing sites, location of RNA degradation signals, etc. For instance, transcription rate of the genes is tuned by transcription factors (TF), which are the most important regulators of gene expression and act as trans elements by binding to promoters and different DNA segments (Chen and Rajewsky 2007). Accordingly, they are one of the key factors in development, differentiation, cell type maintenance and cell signalling pathways including immune responses (Davidson and Erwin 2006; Fong and Tapscott 2013; Lee and Young 2013; Singh et al. 2014).

Besides transcription factors, enhancers and silencers are cis acting transcription regulatory regions, which have fundamental role in gene expression by recruiting and binding to gene controlling machinery. Transcription enhancers and silencers provide binding sites for DNA binding domain of TFs (Berman et al. 2002; Shrinivas et al. 2019). These elements are composed of related motifs as clusters and could be detected by searching for the related clusters in the DNA sequence (Frith et al. 2002; Frith et al. 2003; Cha and Zhou 2014). These clusters are scattered inside the genome.

Riboswitches are one example of gene expression regulatory elements which include clusters of functional motifs in their sequence (Mandal and Breaker 2004; Garst et al. 2011). We have previously reported different riboswitches family-specific sequential blocks, scattered inside the mRNA untranslated regions (UTRs) with varying locations from each other using a block finding algorithm. (Golabi et al. 2018; Golabi et al. 2020b; Golabi et al. 2020a). Similar to other conserved sequence motifs, the riboswitches’ family-specific conserved blocks are small DNA fragments and have almost constant size with some alterations, distributed with varying distances from each other inside the DNA. Therefore, development of new tools for detection of groups or clusters of multiple segments with altering features will be a great help. The found clusters could be binding locations for TFs, riboswitches, suppressors and enhancers, etc., which eventually contributes in determining functional elements and genome structure.

In this paper, we describe a new algorithm and the related web application called FMSClusterFinder, developed for searching clusters of specified functional blocks, with user-defined conditions, in DNA, RNA or protein sequences.

## Materials and Methods

### Cluster finding strategy

This algorithm utilizes the query blocks’ sequences, by considering all the outlined conditions, to determine the maximal clusters present in the subject sequence which fulfil the defined conditions. If the features of the cluster fall within the specified thresholds, the found motifs are displayed as a consequent cluster.

FMSClusterFinder is implemented as a web-based software interface. The core program is written in Python. Input and output stream of the data are handled by PHP. Additionally, the website has been designed using HTML and CSS. Also, JavaScript is used to add more functionality to the website. FMSClusterFinder is designed to help researchers to investigate genomic contents based on their functional regions. FMSClusterFinder can be accessed freely at http://fmsclusterfinder.fmsbiog.com.

### Terms and parameters

- Subject Sequence: the input sequence (DNA, RNA or protein) entered by the user in FASTA format.
- Query Blocks Sequence: one or more motifs which form the cluster with defined minimum and maximum distances between every two proceeding blocks. The query block sequence can be entered in standard and degenerate patterns or have multiple formats.
- Distance to the Next Block: the number of nucleotides/amino acids after the end of one block to the start of the next block is considered as the distance to the next block.
- Similarity Percent: the minimum percent of similarity for any block with the detected motif inside the subject sequence.
- Minimum number of blocks to be present: threshold of the number of necessary blocks to form the proposed cluster.

### Algorithm evaluation

In other to investigate the functionality of FMSClusterFinder program, we worked with a large number of subject sequences in various conditions. Herein, we report the application of the developed program on 120,000 nucleotides (1-120,000) of *Janibacter* sp. HTCC2649 1099316001545 whole genome shotgun sequence, with Genebank accession number of AAMN01000002.1, as a subject sequence example. The sequence contains two SAM-IV riboswitches in its 4575-4686 and 91115-91229 locations according to Rfam (Kalvari et al. 2017; Kalvari et al. 2018). In a previous study analysing SAM-IV riboswitch family blocks (with 3 and more nucleotides), we identified 5 blocks consisting of UCA, GAG, CAG, GCUGG and CGGCAACC motifs (Golabi et al. 2020a). In order to find the possible presence of SAM-IV five motifs cluster inside the *Janibacter* sp. whole genome, we entered the sequences of query blocks and set the required parameters as needed.

Moreover, to examine the accuracy of the program performance, the subject sequence was modified as follows: a) one or more additional clusters were inserted into the sequence, b) one or more clusters were removed from the sequence, c) one block from one of the detected clusters inside the subject sequence was removed, d) the nucleotide sequence of one of the detected blocks inside the subject sequence was changed, and e) the order of the blocks inside the detected clusters was changed.

## Results

### The algorithm features and functionality

The developed “Cluster Finder” algorithm is able to detect clusters of sequential blocks (query blocks) inside a DNA, RNA and protein sequence of interest. This algorithm has the following features and potentials in finding the proposed clusters:

- The user can input a cluster of sequential blocks to be searched for inside the subject sequence.
- Any cluster consists of multiple sequential blocks. Searching for one single block is also possible.
- Any query block’s sequence can be defined in standard (e.g. AAGUC) or degenerate (e.g. CAMT) format.
- In addition to degenerate format, multiple formats for the blocks can also be specified. For instance, for a block composed of 5 nucleotides, the user can state three alternatives as ACCGU-ACGCU-ACGGU meaning that the block can be in one of these 3 formats. The specified multiple formats are different from degenerate format of ACSSU. ACSSU refers to 4 blocks of ACCCU, ACCGU, ACGCU and ACGGU, while the mentioned multiple formats include only three formats. Therefore, the algorithm provides the possibility of defining the blocks in standard, degenerate and multiple formats.
- The desired distance between the blocks could be either a constant or varying number of nucleotides. For instance, one can ask to search for 3 blocks with the following distances:
  - distance between blocks 1 and 2: 15 nucleotides
  - distance between blocks 2 and 3: 14-22 nucleotides.
  So, the detected cluster should be with the following arrangement: **Block 1 →** (15 nucleotides) **→ Block 2 →** (14-22 nucleotides) **→ Block 3** Therefore, the presence of the three blocks located within the defined ranges will be considered as a cluster.
- The boxes of “query blocks sequence” and “distance to the next block” are draggable and the order of the blocks could be readjusted.
- The minimum acceptable number of blocks out of total number of query blocks can be verified. For instance, the user can enter 5 query blocks and then define that presence of at least 3 blocks to form the cluster is acceptable.
- The similarity percentage (similarity percent) for each query block can also be stated. For instance, for the block of “ACG”, if the user defines 66% as the similarity percent, it means the presence of at least 66% of the sequence nucleotides of ‘ACG’ is acceptable. Therefore, the presence of 2 nucleotides out of 3 nucleotides within the sequence will be acceptable and consequently ‘AC’, ‘CG’ and ‘ANG’ segments inside the query sequence will be considered as the block. The similarity percent for each block is defined individually.
- Moreover, this algorithm can search for many motifs individually, without considering them as a cluster. In this case the blocks are entered in any order and their minimum and maximum distances should be set to zero and the minimum number of the acceptable blocks is stated as 1.
- A graphical abstract of the procedure is presented in Supplementary File S1.

### An example

As mentioned, we used FMSClusterFinder to detect the presence of SAM-IV five motifs cluster inside the *Janibacter* sp. whole genome, we entered the sequences of 5 query blocks including UCA, GAG, CAG, GCUGG and CGGCAACC motifs and set similarity percent for all blocks as 100, meaning that the sequence of the found blocks inside the subject sequence should be 100% identical with the query blocks sequence. The minimum and maximum distances between every two adjacent blocks were defined as: blocks 1 and 2, 0-5 nucleotides; blocks 2 and 3, 0-7 nucleotides; blocks 3 and 4, 12-20 nucleotides; blocks 4 and 5, 0-5 nucleotides. As mentioned earlier, the distance between two blocks in FMSClusterFinder is measured between the last nucleotide of the one block and the first nucleotide of the next block. The minimum number of the blocks to form a cluster is also defined as 5, meaning that all the query blocks should be present within the specified distances so that the cluster can be acceptable as a SAM-IV riboswitch related cluster.

Fig. 1 shows the FMSClusterFinder web application display after insertion of input data including: *Janibacter* sp. whole genome as the subject sequence, 5 query blocks, minimum and maximum distances of any two proceeding blocks, the similarity percent and minimum number of the blocks thresholds.

**Fig.1.**
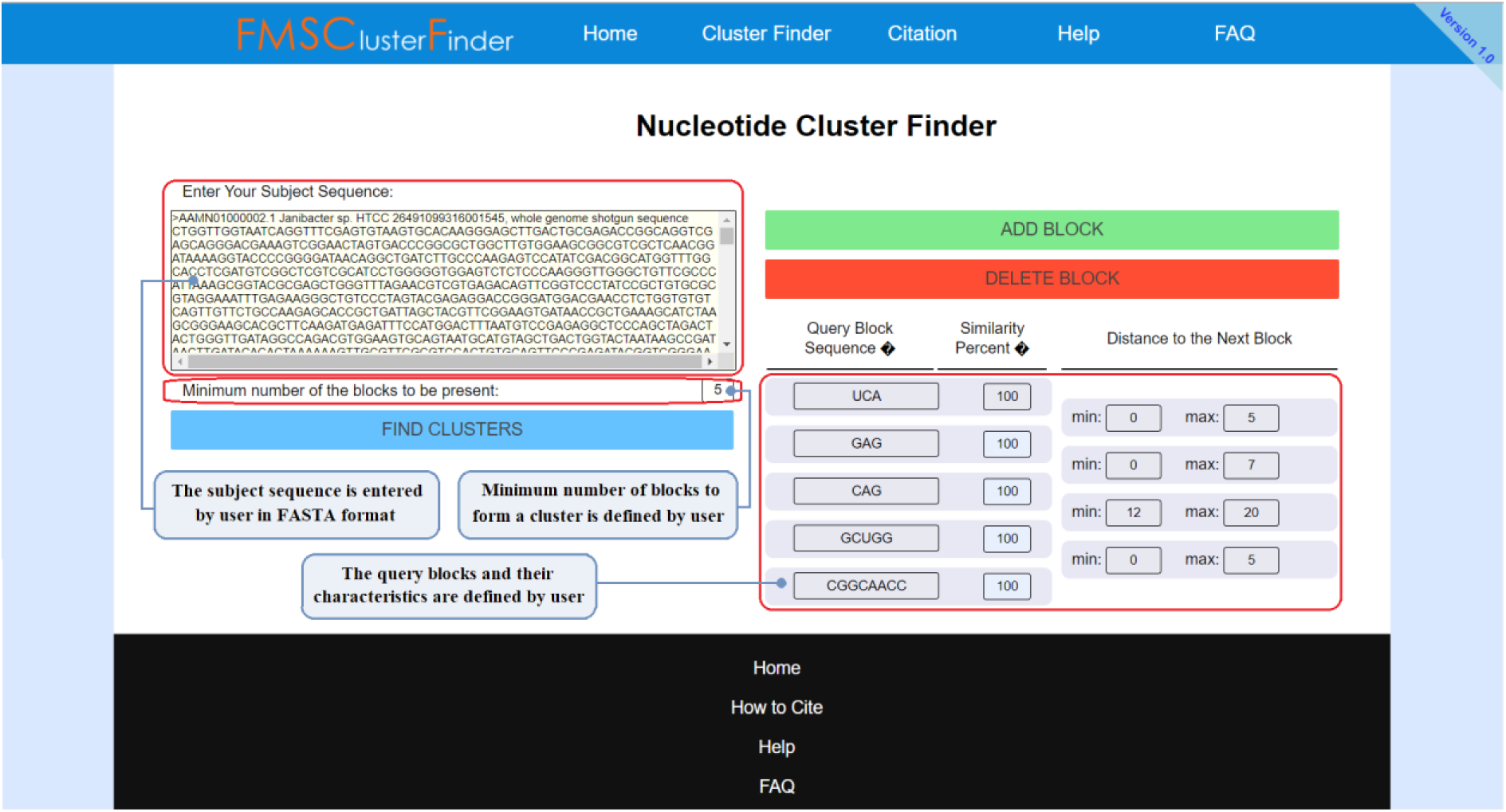
FMSClusterFinder input page.

As seen in the results page (Fig. 2 and Supplementary File S2), the algorithm confirmed the presence of three clusters. The found clusters contained all 5 entered query blocks within the pre-defined range of distance (5 of 5 blocks found). The first detected cluster starts at location 4577 and ends at 4619. This corresponds to the SAM-IV riboswitch related to locations 4575-4686. The second detected cluster starts at location 91117 and ends at 91159, which corresponds to the SAM-IV riboswitch related to locations 91115-91229. These results confirm the performance of the algorithm. As seen, the FMSClusterFinder outputs are in accordance with Rfam data (20, 21). The third cluster (containing/composing SAM-IV riboswitch blocks with three and longer nucleotides as defined in input data) is also detected at locations 104084 to 104126. By clicking on each result, additional information related to the cluster such as the exact locations of the detected motifs is shown. The results page also shows the number of finalized clusters and the execution time.

**Fig.2.**
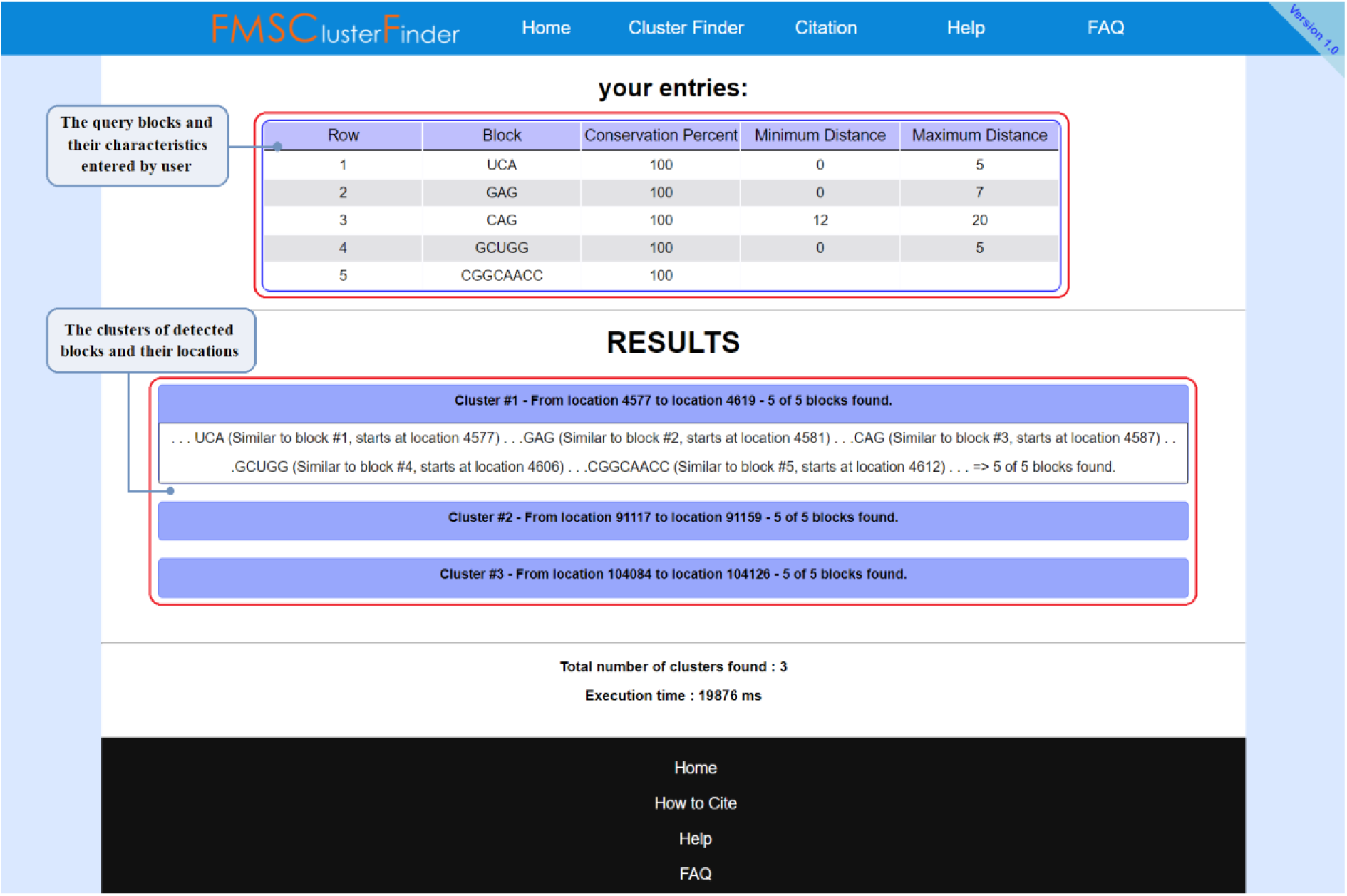
FMSClusterFinder results page.

Moreover, manipulation of the subject sequence by addition or removal of one or more clusters inside and the subject sequence showed that the number of the detected clusters is increased and decreased by insertion or deletion of the clusters, respectively. As explained in materials and methods section, the program performance was also evaluated by removing the blocks, changing one block’s nucleotide sequence and also altering the order of the blocks inside the subject sequence. The results demonstrated that the number of the detected clusters was decreased according to any of the mentioned modifications accordingly. All the observations confirmed the accuracy of the developed program function.

## Discussion

The discovery of genomic functional features has raised interest in prediction and determination of functional sites in DNA and RNA sequences. Since most of these functional areas consist of clusters of specific motifs, bioinformatics tools to detect such clusters based on their characteristics are needed.

We designed an algorithm for detection of clusters consisting various arrangements of query blocks in subject sequences of DNA, RNA or protein. Any number of succeeding blocks can be defined by the user with any desired lengths, conservation (similarity) percentage and the blocks’ sequence could be addressed in standard, degenerate or multiple formats. Meanwhile, various blocks could be searched independently as individual motifs or in association with each other. When a cluster of blocks is searched, the minimum and maximum distances between each two following blocks in the cluster can be stated. Furthermore, a minimum number of occurrences for the blocks out of all entered query blocks can be specified. The FMSClusterFinder web application representing the designed algorithm, is a user-friendly application with rapid execution. It is also free to use and requires no sign-in, download or installation.

## Acknowledgment

The authors thank Professor Iftikhar J. Kullo and Dr. Mojtaba Parvizi (Mayo Clinic, Rochester, Minnesota, US) for their assistance in editing the manuscript.

## Compliance with ethical standards

### Ethical approval

This article does not contain any studies with human participants or animals performed by any of the authors.

## References

Berman BP, Nibu Y, Pfeiffer BD et al. (2002) Exploiting transcription factor binding site clustering to identify cis-regulatory modules involved in pattern formation in the Drosophila genome. Proceedings of the National Academy of Sciences of the United States of America 99:757–762

Cha M, Zhou Q (2014) Detecting clustering and ordering binding patterns among transcription factors via point process models. Bioinformatics (Oxford, England) 30:2263–2271

Chen K, Rajewsky N (2007) The evolution of gene regulation by transcription factors and microRNAs. Nat Rev Genet 8:93–103

Davidson EH, Erwin DH (2006) Gene regulatory networks and the evolution of animal body plans. Science 311:796–800

Fong AP, Tapscott SJ (2013) Skeletal muscle programming and re-programming. Curr Opin Genet Dev 23:568–573

Frith MC, Li MC, Weng Z (2003) Cluster-Buster: Finding dense clusters of motifs in DNA sequences. Nucleic Acids Res 31:3666–3668

Frith MC, Spouge JL, Hansen U, Weng Z (2002) Statistical significance of clusters of motifs represented by position specific scoring matrices in nucleotide sequences. Nucleic acids research 30:3214–3224

Garst AD, Edwards AL, Batey RT (2011) Riboswitches: structures and mechanisms. Cold Spring Harb Perspect Biol 3

Golabi F, Shamsi M, Sedaaghi MH, Barzegar A, Hejazi MS (2018) Development of a new sequential block finding strategy for detection of conserved sequences in riboswitches. BioImpacts 8:13–22

Golabi F, Shamsi M, Sedaaghi MH, Barzegar A, Hejazi MS (2020a) Classification of Riboswitch Families Using Block Location-Based Feature Extraction (BLBFE) Method. Advanced Pharmaceutical Bulletin 10:97–105

Golabi F, Shamsi M, Sedaaghi MH, Barzegar A, Hejazi MS (2020b) Development of a new oligonucleotide block location-based feature extraction (BLBFE) method for the classification of riboswitches. Molecular Genetics and Genomics 295:525–534

Ian Dunham, Anshul Kundaje, Shelley F Aldred et al. (2012) An integrated encyclopedia of DNA elements in the human genome. Nature 489:57–74

Kalvari I, Argasinska J, Quinones-Olvera N et al. (2017) Rfam 13.0: shifting to a genome-centric resource for non-coding RNA families. Nucleic Acids Research 46:D335–D342

Kalvari I, Nawrocki EP, Argasinska J et al. (2018) Non-Coding RNA Analysis Using the Rfam Database. Curr Protoc Bioinformatics 62:e51

Lee TI, Young RA (2013) Transcriptional regulation and its misregulation in disease. Cell 152:1237–1251

Mandal M, Breaker RR (2004) Gene regulation by riboswitches. Nat Rev Mol Cell Biol 5:451–463

Shrinivas K, Sabari BR, Coffey EL et al. (2019) Enhancer Features that Drive Formation of Transcriptional Condensates. Mol Cell 75:549–561.e547

Singh H, Khan AA, Dinner AR (2014) Gene regulatory networks in the immune system. Trends Immunol 35:211–218

Yue F, Cheng Y, Breschi A et al. (2014) A comparative encyclopedia of DNA elements in the mouse genome. Nature 515:355–364

