## Supplementary figures and images for "FMSClusterFinder: A new tool for detection and identification of clusters of sequential motifs with varying characteristics inside genomic sequences"

### Supplementary File S1

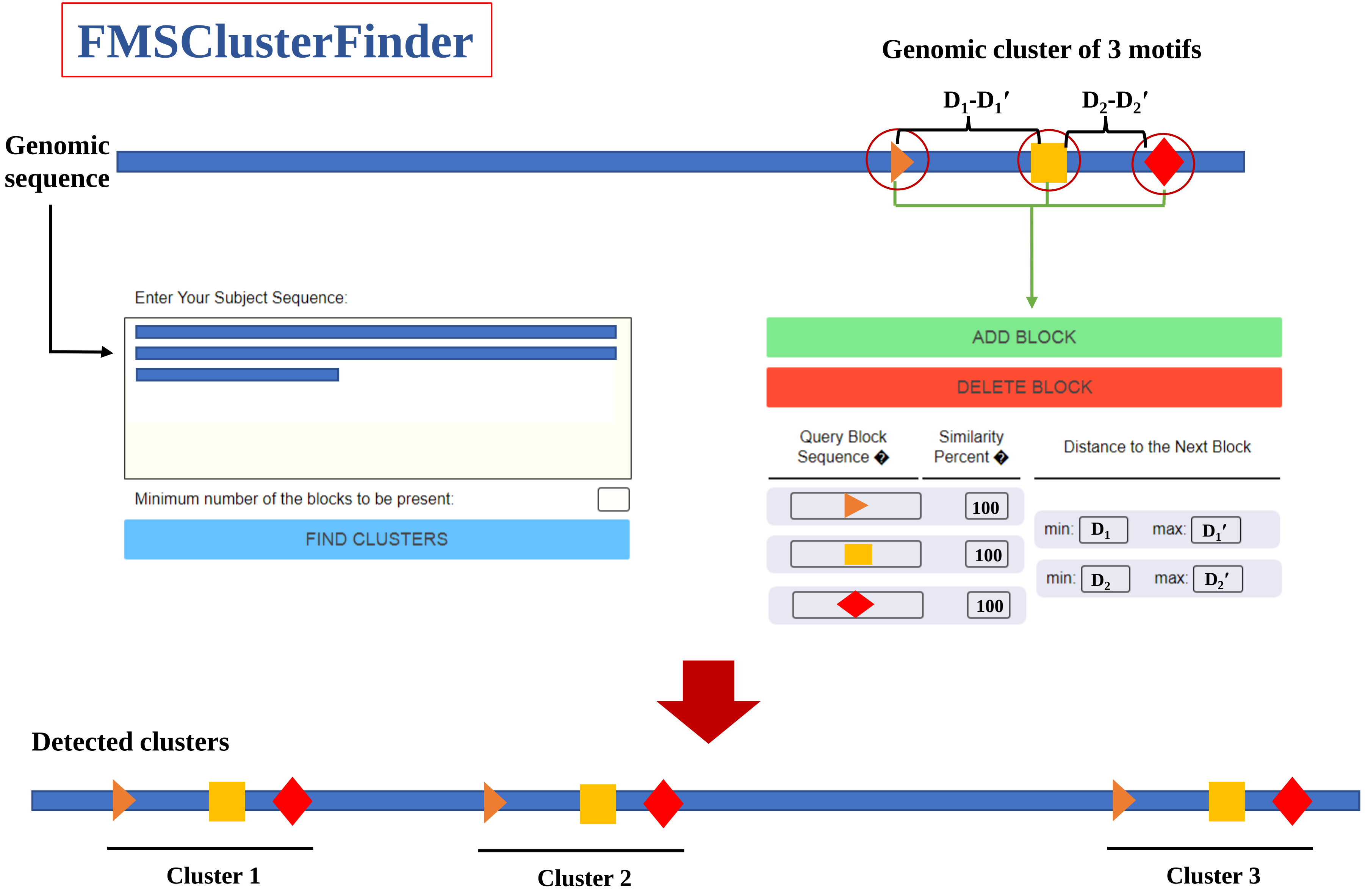
