## Supplementary File S2 for "FMSClusterFinder: A new tool for detection and identification of clusters of sequential motifs with varying characteristics inside genomic sequences"

**Cluster 1 (5 blocks): starts at 4577**

CAACCGGTGGGCGCCCAAGAGCCTGCTCCTGCTCCCGATGGGTGTCCTTGCCCTCGGCGGTGTCGTCGTGAT
CGGCATGGGGCTACTCGAAGGGGGCGACGACCGAGCCCTGAAGATCGTGGGCGCGCTCCTCGCAATCGCGT
CCCTCGTCATCACCGGAGCCGCCAACGTCCCGATCAACAACAAGGTCGACGCGCTCACCGCAGGGTCGGAAG
CGGAAGGCGAGTGGGCGGCATACGCCGCGGTGTGGTGCCGATGGAACCACCTCCGCACCGTGACGAGTCTC
GCCTCAGCCGTGCTCCTCGCGGTGGCCGCCGCGCGGACCTGACCGTCTCGGGTGGTGGACGGGGGTCCCC
TGCACGTGCGACCGAGCCATAGACTCCACAACCAAGG**TCA**C**GAG**TAC**CAG**TGTTCAGCCCCGGCTT**GCTGG**T
**CGGCAACC**CTCCTCCGCGGTGGGGTGCTCCGGGTGACGACCTGGTCGGCTGCAGCAACGCAGTTGGCAAGC
GCGGACTCCCGAGCACCTTCGAGGGTCCAACGACTCTCCAGGGAGCACCGGATGTCCATCGCCAGCTCGATTC
CTCACACCTTCACCCGTGAACGTCGCCAGCCGTGGCTGCGCTCCGTCCCCGCACCGGCCGGGCCCGCCACCCC
GAGCGCGCCCAGCGACGACAACGTCTCGCATGCGGACGGCATACCCACCGACGTTGTGGACACCAAGGTGCC
GGCGGTCGTTGCCGCACCCAAGGTCCCGACGTGTCACGGCACCTTCGTCGAGTACGCGAACCTCGACCACGG
CGCCTCGACCCCGGCACTGGCCAGCGTGGTCGCGGCCGTTGACAAGGCGACCCAGACCTACTCGAGCGTCCA
CCGC

**Cluster 2 (5 blocks): starts at 91117**

CCCCTGTCGAACGCGCGGCGCAGGATCGCCTGCTGACGGTCGAGGGGCACGTCGTCACCGAAGTTGTGCCAC
AGGCCGAGAGAGATTGCAGGGAGGTCGAGACCGGACCGCCCGGCCCGGCGATACGGCATACCCGCGCGGTA
GCGGCCAGGGGCCGCCTTGTACTCATCGACGTGGAAAGTCATGTGACTCATCATGCCTGCCTATGGCGGTCCA
CGATCAGAGACACCGCTCCCGCGGGCAGGGGCGGCGGCGTTAACGTTTCAACCACAGG**TCA**T**GAG**TGC**CAG**
CGACAAGCCCCGGCTC**GCTGG**C**CGGCAACC**CTCCAACCGCGGTGGGGTGCCCCGGGTGAAGACCAGGCGGA
GCGTCGACCGGCGCCCGCAAGCACGGCATACAAGGAGCTTTTGCGATGGCTGACGACGCAACTTCTGGCACC
CCGAACTGGTCGTTCGAGACCCGGCAGATCCACGCCGGACAGGTGCCCGACCCGACGACCGGCGCCCGCGCG
CTACCGATCTACCAGACGACCTCCTACGTCTTCAAGGACACCGAGCAGGCGGCCAACCTGTTCGCGCTCAAGG
AGTTCGGCAACATCTACACGCGCATCATGAACCCGACCCAGGACGTCGTGGAGCAACGCATCGCCTCGCTGG
AAGGCGGCGTCGGCGCTTTGCTCGTGGCCTCGGGTCAGGCCGCGGAGACCATCGCGATCCTCAACATCGCCG
AGGCCGGCGACCACATCGTCGCGTCCCCGTCGCTCTACGGCGGCACGTTCAACCTGCTCAAGCACACGCTGCC

**Cluster 3 (5 blocks): starts at 104084**

GAGGCGGCAGCGATCTGCACGATCGCCGCGTGCGTCGCCTGGGGCATCGGCAACGGCTCCACGGCGGGTGA
TGCGGTGACCCGTGCGCTGCGCGTCGACCCGGACTATCGCCTTGCGAGGTACCTCGGCCGGATGATCGACCA
CCAGATGCGCCCTCGCCACGCCTGGGCCGATGTCGCCGCCTGAGCGCGTGGCTGGAGGTCCCTCTCGGCTGT
CGCACTCTCACGCTGCGGACAGGGTGCCGTGGAAGGGGTCACCGGTTCTAGACTCGTGCCCAGG**TCA**T**GAG**T
CC**CAG**CGACAAGCCCCGGCTT**GCTGG**G**CGGCAACC**CTCCTCGCGGTGGGGTGCCCCGGGTGAAGACACGGC
CCTTCCGGTATCGGTGGGGCAAGCGCGATCCGAGGAGCACTCCGATGACCATCACCGACCTCCCGACAGCGT
CGTTCTGGCGCCCGGGCGATCACCCGGGCGCGCGCCAGTTCGTCCAGGTCGGCCCCGTCGCGCTCGAGCGCG
GTGAATCCCTGCCCGAGGTCACCGTGGCCTACGAGACGTGGGGGACGCTCAACGCCGCCCGCGACAACGCG
GTGCTCGTCGAACACGCCCTCACCGGCGACGCCCACGTCGAGGGCCAGGCCGGTCCCGGCCACGCCACGCCC
